# A Model Selection Approach to Hierarchical Shape Clustering with an Application to Cell Shapes

**DOI:** 10.1101/2020.04.29.067892

**Authors:** Mina Mirshahi, Vahid Partovi-Nia, Masoud Asgharian

**Affiliations:** Department of Mathematics and Industrial Engineering, Polytechnique Montreal, Canada; Department of Mathematics and Statistics, McGill University, Montreal, QC, Canada

## Abstract

Shape is an important phenotype of living species that contain different environmental and genetic information. Clustering living cells using their shape information can provide a preliminary guide to their functionality and evolution. Hierarchical clustering and dendrograms, as a visualization tool for hierarchical clustering, are commonly used by practitioners for classification and clustering. The existing hierarchical shape clustering methods are distance based. Such methods often lack a proper statistical foundation to allow for making inference on important parameters such as the number of clusters, often of prime interest to practitioners. We take a model selection perspective to clustering and propose a shape clustering method through linear models defined on Spherical Harmonics expansions of shapes. We introduce a BIC-type criterion, called CLUSBIC, and study consistency of the criterion. Special attention is paid to the notions of over- and under-specified models, important in studying model selection criteria and naturally defined in model selection literature. These notions do not automatically extend to shape clustering when a model selection perspective is adopted for clustering. To this end we take a novel approach using hypothesis testing. We apply our proposed criterion to cluster a set of real 3D images from HeLa cell line.

## Introduction

Shape modelling plays an important role in medical imaging and computer vision [1, 2]. There is, in particular, a vast literature on cell shape analysis where the shape can often provide important information about the functionality of the cell ([3, 4, 5, 6, 7]). Most of such analysis employ techniques that often lack a statistical error component. Variability among shapes has been studied using different approaches: active shape models [8], computational anatomy [9], planar shape analysis [10, 11], etc.

Hierarchical clustering is one of the most commonly used methods by practitioners to find unknown patterns in data [12]. There are several reasons why a tree based method is preferable: i) the closest objects merge earlier, which reflects the evolution as the hierarchies are built up ii) provides a visual guide through a binary tree, called dendrogram iii) For a chosen number of clusters, it is only required to cut the dendrogram to achieve the corresponding clustering iv) allows to choose the number of clusters visually so that corresponding clustering is meaningful to an expert.

The existing hierarchical shape clustering methods are distance based and often lack a proper statistical foundation for making statistical inference. Borrowing ideas from model selection and testing statistical hypothesis, we take a different perspective to hierarchical clustering. A statistical shape model built on a set of image data, using a common coordinate system, can be regarded as a random continuous curve. A 3D shape can then be represented as a linear function of some basis functions. Possible noises and fluctuations can be easily included in the linear model. One can therefore use the likelihood function to devise a shape clustering algorithm using a convenient metric. The main advantage of such approach is to acquire the probability distribution of the fitted function and to distinguish between different shapes using their probability distributions.

Having modelled shapes using linear models, clustering can be performed on the estimated coefficients of the basis functions; similar shapes are expected to have similar coefficients. The problem of assigning two shapes to the same cluster will then become a hypothesis testing problem. Having this formulation, choosing the optimal number of clusters can be treated as a model selection problem. To this end, we use BIC and devise a criterion, called CLUSBIC, whose consistency is established.

The rest of this manuscript is organized as follows. In Section we discuss statistical modeling of shapes in three-dimensions using spherical harmonics. In Section 1 we propose a new Bayesian information criterion, called CLUSBIC, for clustering shapes taking into account the likelihood function of their fitted models, and establish consistency of the proposed criterion. In Section 2 we apply our proposed method to 3D images from HeLa cell line captured by a laser-scanning 240 microscope. Section 3 is the conclusion. Proofs of the main results are given in the appendices, while the proofs of other auxiliary results and the lemmas are documented in the supplementary materials.

sectionShape Modelling A three-dimensional (3D) shape descriptor is highly beneficial in many fields such as biometrics, biomedical imaging, and computer vision. Double Fourier series and spherical harmonics are widely used for representing 3D objects. Here, we discuss spherical harmonics for 3D shape modelling. Spherical harmonics are a natural and convenient choice of basis functions for representing any twice differentiable spherical function.

Let *x, y* and *z* denote the Cartesian object space coordinates, and *θ, ϕ*, and *r* the spherical parameter space coordinates, where *θ* is taken as the polar (colatitudinal) coordinate with *θ* ∈ [0, *π*], and *ϕ* as azimuthal (longitudinal) coordinate with *ϕ* ∈ [0, 2*π*]. Spherical harmonics is somehow equivalent to a 3D extension of the Fourier series, which models r as a function of *θ* and *ϕ*. The real basis for spherical harmonics 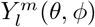 of degree *l* and order *m* is defined as:

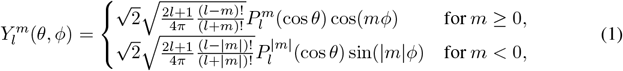

where *l* and *m* are integers with |*m*| ≤ *l*, and the associated Legendre polynomial 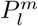 [13].

Given a spherical function *r*(*θ, ϕ*) and a specified maximum degree *L*_max_, one can write *r*(*θ, ϕ*) as a linear expansion of 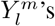 with possibly some measurement errors and find the coefficient of fit through the method of least squares. That is,

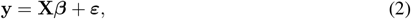

where

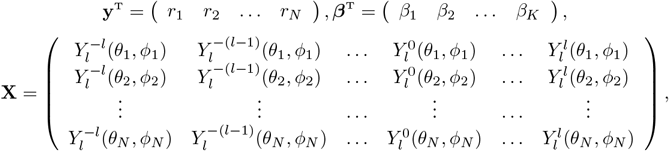

where *N* is the number of observations, |*m*| ≤ *l* ≤ *L*_max_, and *K* = (*L*_max_ + 1)^2^ is the number of parameters. The quality of fit improves as *L*_max_ increases, i.e., the larger the number of expansion terms.

The model (2) is only suitable for surface modeling of stellar shapes. A different parametric form for surface modeling, regardless of the type of shape, is suggested by [14, 15]. This parametric form gives us three explicit functions defining the surface of shape as *x_s_* (*θ, ϕ*), *y_s_* (*θ, ϕ*), and *z_s_* (*θ, ϕ*), where each of the coordinates is modeled as a function of spherical harmonic bases. Accordingly, the following three linear models are generated,

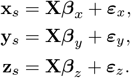

The set of expansion coefficients (*β_x_, β_y_, β_z_*) defines the shape completely. Assuming **x**_*s*_, **y**_*s*_ and **z**_*s*_ to be independent of each other, the above three models are equivalent to the following model

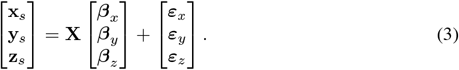

## 1 Shape Clustering

Having characterised shapes by functional forms, we can cluster shapes according to their estimated coefficients. Clustering shapes can be achieved simply by computing the Euclidean distance over their parameters of the models in equation (2). However, this heuristic method may not lead to proper clusters, so we aim to develop a methodology using fitted models. To this end, we present a likelihood-based approach for clustering shapes in this section. From the Bayesian point of view, the clustering procedure is enhanced by contemplating the distribution of parameters in equation (2). We assume some prior distributions over parameters of the fit *β*, and calculate the marginal distribution of the model by integrating the likelihood with respect to the assumed prior. In this approach, the hierarchy of clusters is built up based on the marginal likelihoods [16, 17, 18, 19]. Agglomerative hierarchical clustering is used to establish the dendrogram in which curves are merged as long as the merge improves the marginal likelihoods, as discussed in [20].

This section is organized as follows. First, a brief sketch of likelihood calculation with classical assumptions is provided in Subsection 1.1. In Subsection 1.2, we introduce a new Bayesian information criterion for clustering curves, called CLUSBIC. In Subsection 1.3 the consistency of the CLUSBIC is proved.

### 1.1 Linear Models

We consider the following linear model in polar coordinates

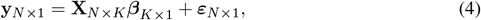

where *N* and *K* indicate the number of observations and the attributes respectively. We assume *ε* is distributed according to the Gaussian distribution with mean **0**_*N* × 1_ and covariance matrix *σ*^2^**I**_N_, i.e. 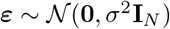, where **I**_*N*_ is the identity matrix of size *N*.

In case of spherical harmonic expansions, equation (3), we assume that the error terms *ε_x_, ε_y_*, and *ε_z_* are independent of each other and each follows the Gaussian distribution 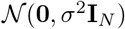. Subsequently, the vector

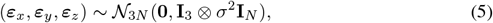

where the symbol ⊗ is the Kronecker product.

Given *D* distinct shapes, the model associated with this set of shapes is

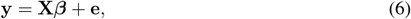

where

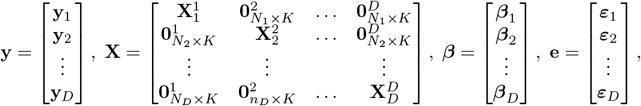

We explain the methodology using the notation for equation (4). Similar methodology applies using equation (3), taking into account the property in equation (5).

Suppose **d** = (*d*_1_, *d*_2_,…, *d_D_*) is a grouping vector, e.g. **d** = (1, 1,…, 1) assigns all *D* shapes to one group and **d** = (1, 2,…, *D*) assigns each shape to a singleton. The likelihood function for the model (6) given the grouping vector **d** is,

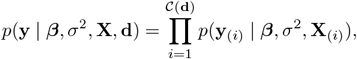

where 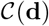 denotes the number of unique elements in **d** (the number of clusters) *N*_(*i*)_, while **Y**_(*j*)_ and **X**_(*i*)_ represent, respectively, the number of observations, vector of response, and matrix of covariates after combining the clusters. The vector of unknown parameters is denoted by *β*.

For the sake of simplicity, we propose conjugate priors [21]. To begin with, *σ*^2^ is assumed to be known. The standard conjugate prior imposed on *β*, conditional on *σ*^2^, is 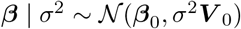, where *β*_0_, and ***V***_0_ are the prior mean and prior covariance matrix for β respectively. For a detailed discussion of Bayesian methods see [22]. The marginal likelihood of **y** can be computed as follows:

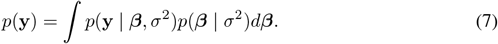

In case of conjugate priors, the model appears as the multivariate Gaussian distribution with mean **X***β*_0_ and covariance matrix of **I**_*N*_ + **XV**_0_**X**^T^, where 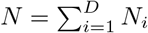 denotes the number of observations. The curves are assigned to a group with maximum *p*(**d** | **y**). By the Bayes theorem,

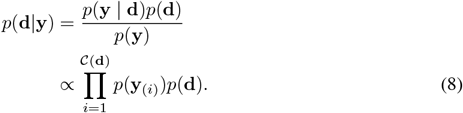

The dendrogram exploits the marginal likelihood to build the hierarchy. It is expected that the marginal likelihood reaches its maximum over a reasonable grouping. Thus, the logical cut-off on the dendrogram is when the marginal likelihood is maximized over the dendrogram.

In order to calculate the marginal likelihood *p*(**y** | *σ*^2^) for each curve, one needs to compute the inverse of the covariance matrix (**I**_*N*_ + **XV**_0_**X**^T^) which is of size *N*, the number of observations. The computation of the inverse has the computational complexity of 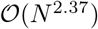 [23] for the best-case scenario. To circumvent the computational complexity involved in computing the inverse of the covariance matrix, we modify the Gaussian model by adding *σ*^2^ to the hierarchy of parameters. Assuming an inverse-gamma distribution for *σ*^2^, transform the Gaussian model to Student’s t-model which does not require any matrix inversion of size *N*, see [24]. If *σ*^2^ is nearly degenerate, i.e, E(*σ*^2^) = *μ*_0_, and Var(*σ*^2^) ≈ 0, this model is, asymptotically, the same as the Gaussian model. The computational complexity of this model is bounded by 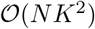 which is a significant improvement over the Gaussian model. The proposed computational trick leads to a marginal likelihood which comes from a distribution with heavier tails than the Gaussian distribution. This trick is particularly helpful in modeling data containing outliers.

As [25] discussed, agglomerative clustering using Student’s t-distribution suffers from some instabilities under the common settings of the hyper-parameters. Here, the hyper-parameters of the inverse-gamma distribution are determined such that it produces a nearly degenerate distribution.

Bayesian clustering according to the *p*(**d** | **y**), equation (8), requires modeling *p*(**y** | **d**) and *p*(**d**). Here, we follow the structure suggested in [20]. The random vector **d** denotes the possible groupings for a set of shapes. Suppose 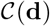 is the total number of clusters at each step and 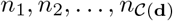 are the total number of shapes in each of the clusters. Suppose that 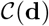 is uniformly distributed over the set {1,2,…, *D*}, where *D* is the total number of shapes and the *n_j_, j* = 1,2,…, *D*, is distributed according to multinomial-Dirichlet with parameter *π*,

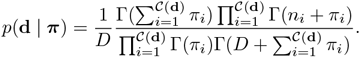

A uniform setting on the parameter vector π leads to

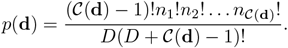

### 1.2 Clustering Bayesian Information Criterion (CLUSBIC)

We introduce a criterion for clustering, based on the marginal likelihoods, called *Clustering Bayesian Information Criterion* (CLUSBIC). CLUSBIC is similar to BIC in nature, but it is designed for the purpose of hierarchical clustering. When all data fall into one cluster, CLUSBIC coincides with BIC.

[26] proposed a simple informative distribution on the coefficient *β* in Gaussian regression models. As a prior on the parameter verctor, given the covariates, he considered 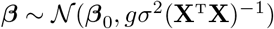, a conjugate Gaussian prior distribution. This covariance matrix is a scaled version of the covariance matrix of the maximum likelihood estimator of *β*. In practice, *β*_0_ can be taken as zero and *g* is an overdispersion parameter to be estimated or manually tuned. The parameter *g* can be set according to various common model selection methods such as AIC, BIC and RIC [27]. [26, 28] suggested the use of the prior distribution on *g* as a fully Bayesian method (see also [29]). Various other methods have been recommended in the literature for finding an optimal value for *g*, such as the empirical Bayes methods [30, 31]. Choosing the number of clusters according to the marginal probability distribution is analogous to using the BIC criterion in a model selection problem if *g* = *N* where *N* is the number of the observations.

Denote by 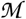 the set of all possible models. The set 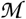 contains all the possible ways that one can assign *D* distinguishable shapes into *D, D* – 1, *D* – 2,…, 1 indistinguishable clusters. The cardinality of 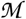 is 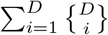 where 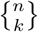 is the Stirling number of the second kind. The sum is equal to the Bell number 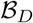.

The testing formulation of model selection is to test the following hypothesis

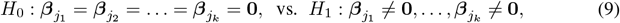

for 1 ≤ *j*_1_ < *j*_2_ < … < *j_k_* ≤ *D*. Clustering, in contrast, is concerned with finding different ways of combining the covariates. Clustering may then be formulated as testing the hypothesis

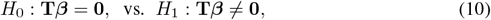

where **T**_*qK*×*DK*_ is the matrix of constraints such that **T***β* = **0** is a set of *q* linear constraints on *β_j_* for 1 ≤ *j* ≤ *D*. The following theorem provides an approximation of the marginal likelihood which we call CLUSBIC.

#### Theorem 1.

*Suppose p*(**y** | *β*, **X**, **d**, *σ*^2^) *is a C*^3^ *function of β. Then as N_i_* → ∞, ∀*i, we have*

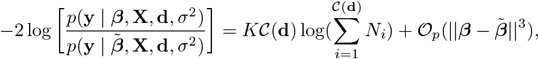

*where* 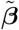 *is the maximum likelihood estimate of β under the null hypothesis, H*_0_: **T***β* = **0**.

*Proof*. The proof is given in Section A of the Appendix.

Theorem 1 shows that for large sample sizes our proposed clustering method using the marginal likelihood has the same premise as Ward’s linkage [32]. Ward’s linkage, Δ(*A, B*), is a popular method in hierarchical clustering which is based on minimizing variance after the merge. Ward’s method merges two clusters A and B that minimizes the sum of squares after the merge.

Using CLUSBIC, two groups, say group *i* and group *j*, are merged together if it leads to a decrease in the CLUSBIC. The CLUSBIC is calculated for different combination of shapes. A pair of groupings that minimizes CLUSBIC is a merging candidate. For example, groups *i* and *j* are merged together in the second level of the hierarchy only if the merge decreases the total CLUSBIC compared to the level before. The following calculation shows that this idea of merging the groups is very much aligned with likelihood maximization.

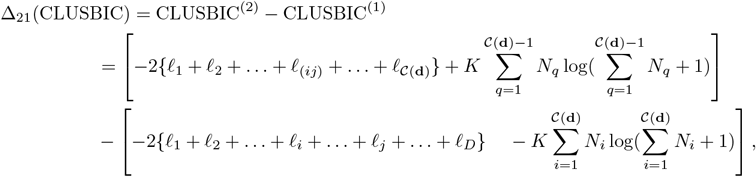

where *ℓ_i_* = log{*p*(**y**_*i*_ | *β_i_*, **X**, *σ*^2^)} and *ℓ*_(*ij*)_ is the log-likelihood after merging group *i* with group *j*. As the total number of observations is fixed at each level of hierarchy, i,e., 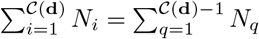

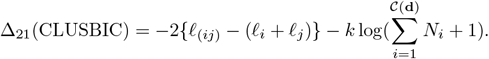

For Gaussian models, Ward’s linkage is closely related to CLUSBIC. Minimizing the CLUSBIC between each two consecutive levels of the dendrogram leads to minimizing the metric, *c_ij_* = *ℓ_i_* + *ℓ_j_* – *ℓ_j_*; *i, j* = 1,2,… *D*. Now, considering the model **y** = **X***β* + *ε*,

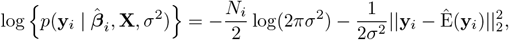

where 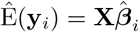. Substituting the above equation in the *c_ij_* metric, we have

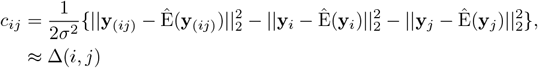

When **X**^T^ **X** is a diagonal matrix, *c_ij_* equals Δ(*i, j*).

### 1.3 Consistency of CLUSBIC

In this section, we show that the CLUSBIC criterion is consistent in choosing the true clustering as *N_i_* → ∞ for 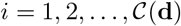. Assuming that there exists a model 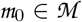 that represents the true clustering, the CLUSBIC, developed in Theorem 1, is said to be a consistent criterion if

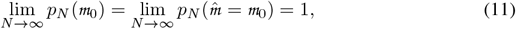

where 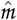 is the model selected by CLUSBIC.

In a regression setting, the space of models 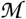 can be partitioned into two sub-spaces of under-specified and over-specified models in a rather straightforward fashion. The space of under-specified models 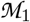 contains all models that mistakenly exclude the attributes of the true models. On the other hand, the space of over-specified models 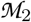 contains all models that include more attributes besides the true model’s attributes. In other words, the sub-spaces 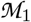 and 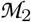 can be effortlessly established considering the presence or the absence of the attributes of the true model. Therefore, more formally, we have

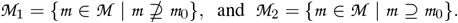

Consequently, for each model that belongs to 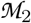, the dimension of the model, *K*, is always greater than the true model.

The definition of under-specified and over-specified models needs to be adjusted for the clustering context. By definition, the over-specified models must include the attributes of the true model. The question that arises here is how the notion of attributes can be interpreted for the clustering problem. Suppose that each column of **X** represents an attribute; more clusters mean more attributes. There is some confusion as to whether the over-specified models should include the clusters of the true model (i.e., smaller number of clusters compared to the true model) or disjoint the clusters of the true model (i.e., having more number of clusters).

Consider the general linear model **y** = **X***β* + **e** in equation 6. Let 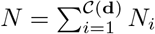 and 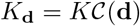 be the total number of observations and the number of parameters in the model respectively. One can approach the problem of defining the model space of 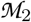 from the hypothesis testing point of view (10). In this approach, the over-specified model is the one that the null hypothesis for that model holds true considering the assumptions on the true model. More formally, we define 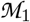 and 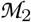 as follows for the clustering problem,

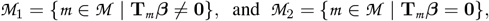

where **T**_*m*_ is the matrix of constraints for the model 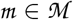. To establish the consistency of CLUSBIC, we need to prove the following statements,

a. lim_*N*→∞ *p_N_*_(*m*) = 0 for 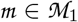.
b. lim_*N*→∞*p_N_*_(*m*) = 0 for 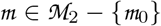.

We in fact prove a stronger version of the first statement (a). We’ll show that

a. lim_*N*→∞_ *N^h^p_N_*(*m*) = 0 for 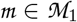, and for any positive *h*.

The consistency of the BIC in model selection has been developed in the literature considering different assumptions, see [33, 34, 35]. Estimating the number of parameters under the null hypothesis in (10) leads to constrained least squares. Consequently,

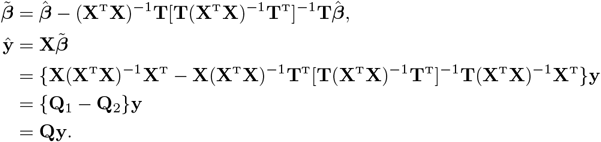

The matrix **Q** is the projection matrix which is symmetric and idempotent.

In the following theorem, we show that CLUSBIC is a consistent clustering measure for any arbitrary distribution on model (4). To prove the consistency of CLUSBIC, we rely on a quadratic approximation to the logarithm of the likelihood suggested by [36]. It can be shown that results can be exact for Gaussian models, see Section B.

#### Theorem 2.

*The* CLUSBIC *is a consistent clustering criterion*.

*Proof*. The proof is given in Section B of the Appendix.

## 2 Application

In this section, we apply our proposed method to the biological cell data obtained from the Murphy lab^1^ ([37]). The database includes 3D images from HeLa cell line captured by a laser-scanning microscope. For this study, we consider images which are labeled as monoclonal antibody against an outer membrane protein of mitochondria. There are fifty data folders each representing the data from a distinct cell. Each folder contains four sub-folders and the data corresponding to the cell and crop image folders are used for this study. The cell folder has various images of a specific cell, taken at different depths called the confocal plane.

To better illustrate our proposed method of clustering, we start by an example of shape clustering in a two-dimensional (2D) space. For this purpose, a single stack, common over all cells, is selected. The chosen stack, thereafter, is segmented using the designed crop image such that there is only one cell per image. To obtain a true representation of a shape from each image, location, scale, and rotational effects should be filtered out. For this purpose, we aligned cells such that their centroids are located in the centre of images. In addition, the cells are rotated in the direction of their main principal component axes to assure a rotation-free analysis. Consequently, the coordinate of pixels on the boundary of the cell is extracted for modeling. We use MATLAB standard methods including segmentation, boundary detection and Savitzky–Golay smoothing filter functions from MATLAB toolboxes [38, 39] for detecting the boundary of the cell. We call the associated line the *oracular boundary* since it, supposedly, represents the true boundary for each cell. Besides, we propose another method for boundary detection which enables us to take into account the associated uncertainty.

For each observation on the oracular boundary, the uncertainty (variance) is calculated based on the data points on the lower and the upper boundaries in polar coordinates. Lower and upper boundaries are treated as a 95% confidence interval and

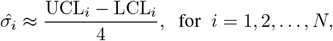

gives an estimation of the standard deviation for point *i*, where UCL_*i*_ and LCL_*i*_ represent the lower boundary and upper boundary values at point *i* respectively. Then the median of the 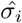 is treated as the common standard deviation to be used for all the computations throughout our clustering algorithm. A Gaussian sample centered around each observation on the oracular boundary with the so-called common variance is generated.

Note that in the case of 2D shape modeling, one can employ basis functions such as Fourier, splines or wavelets and then follow the same procedure for shape clustering as in Section 1. For the sake of readability, in Figure 1, the dendrogram is reported for only ten random cells. Each cell is assigned a number corresponding to its order in the database.

**Fig 1.**
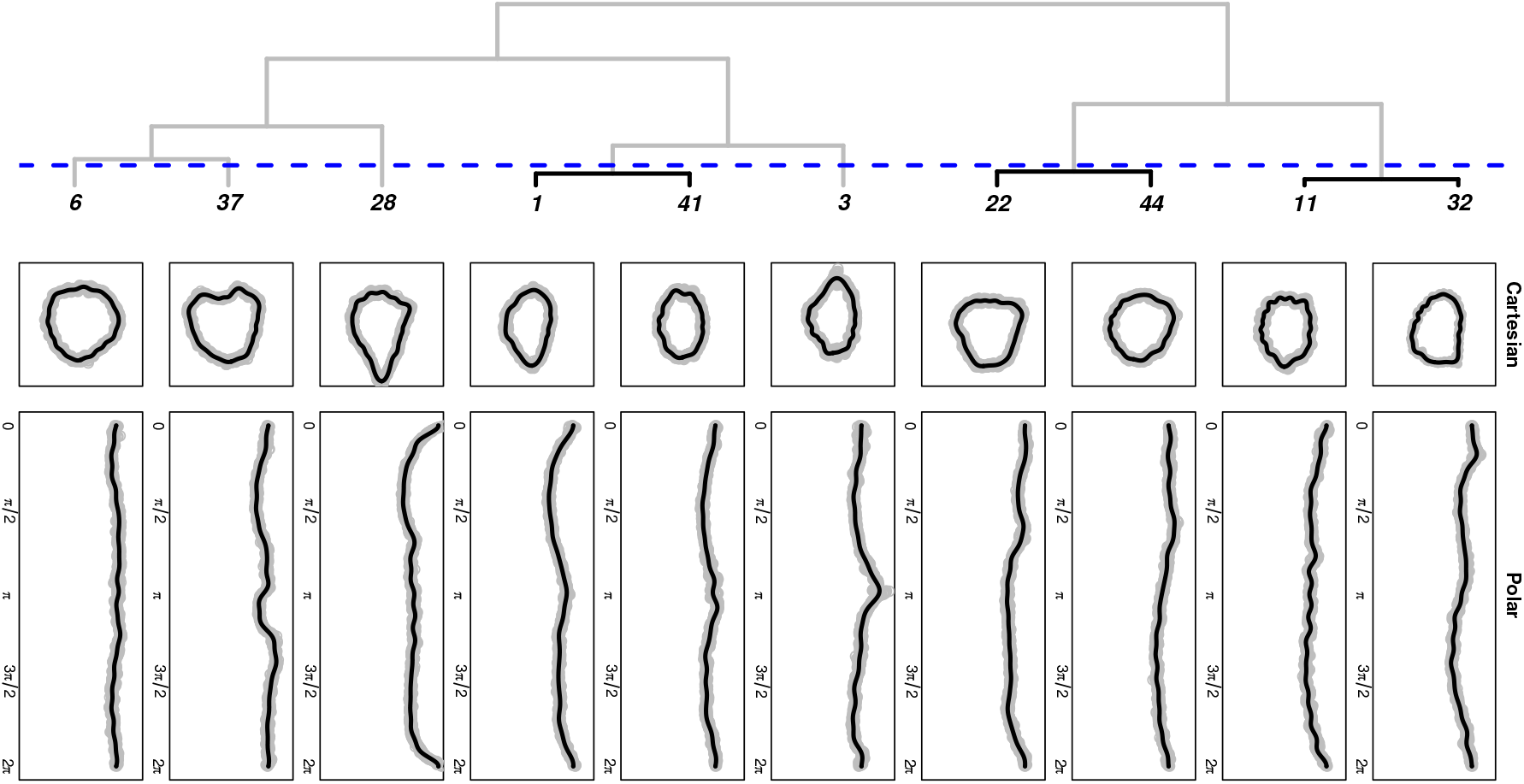
Top panel, dendrogram of the marginal likelihood associated with each cell using Fourier basis functions with *K* = 33 expansion terms. In this example, *D* = 10, *N_i_* = 1000 for *i* = 1, 2,…, 10, *d* = (1, 3, 6, 11, 22, 28, 32, 37, 41, 44) and 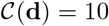. Black lines, in dendrogram, represent the improvement in the marginal likelihood and gray lines depict the deterioration in the marginal likelihood. The dashed blue line indicates the maximum a posteriori cutting point for the dendrogram, d = (1, 3, 6, 11, 22, 28, 11, 37, 1, 22) and 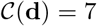. Middle panel, the fitted curves to each of ten random selected cells used in dendrogram in 2D space. The gray points represent the boundary data used in modeling. Bottom panel, the same curves as the middle panel are depicted a in 1D space.

In order to obtain the 3D Cartesian coordinates associated to each voxel on the surface, we do as follows. First, the boundary data for each image stack is extracted separately. Second, the 2D coordinates of each stack are combined all together to create the 3D coordinates of the cell shapes depicted in Figure 2. We then take the image spacing information into acccount.

**Fig 2.**
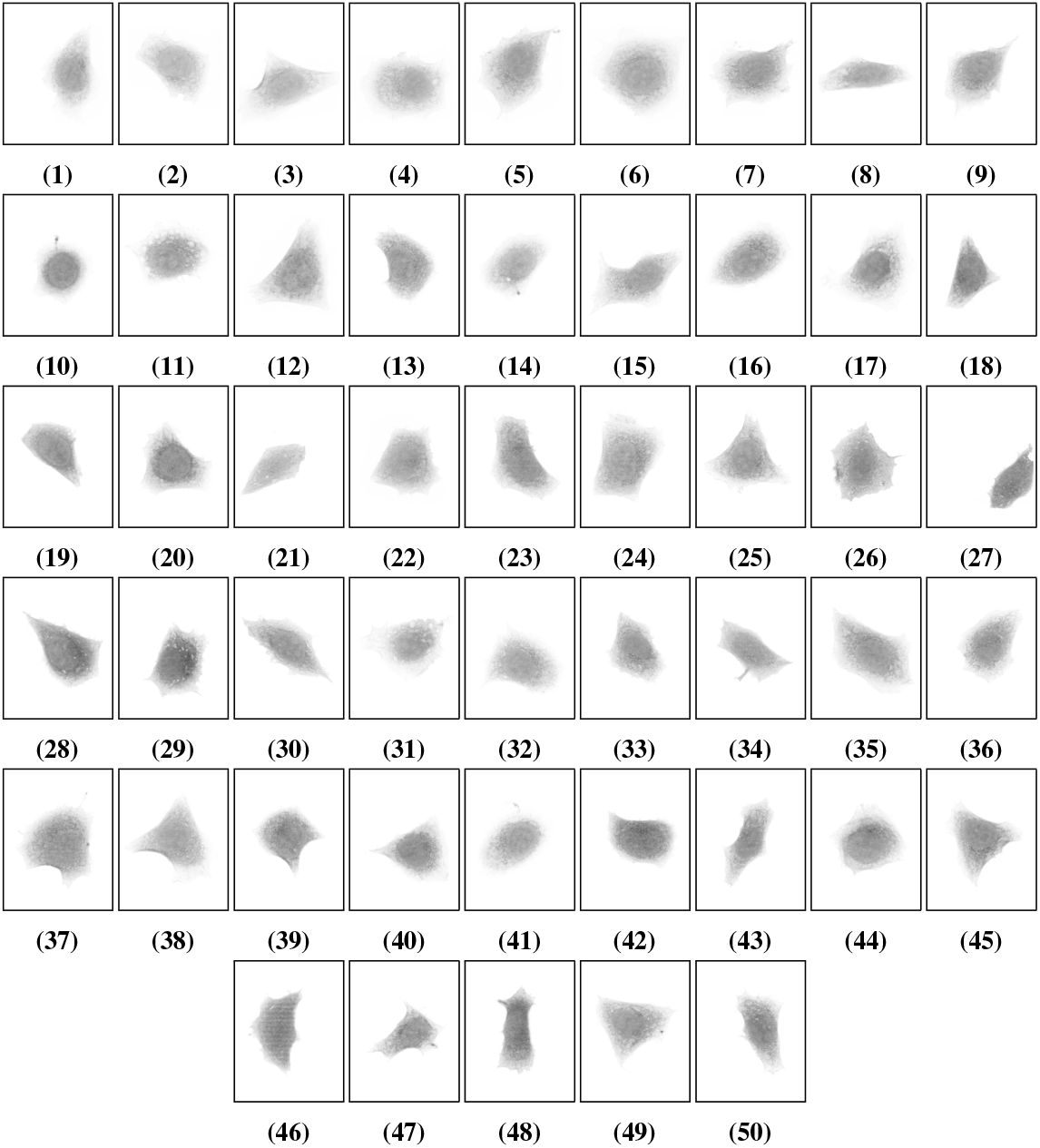
The raw images of fifty cells(D=50) used for clustering throughout this work. The number assigned to each cell matches with its order in the dataset.

For this dataset, the voxel spacing is (0.049μm, 0.049μm, 0.203μm) [40]. It should be noted that the final extracted data must be within a sphere of unit radius to be suitable for modeling using spherical harmonics.

As image data involve noise, the least squares equation (3) is ill posed. A regularization approach can then be taken to estimate the model parameters. See [41] for details and further discussion. The result of clustering for *D* = 10 random cells is reported in Figure 3.

**Fig 3.**
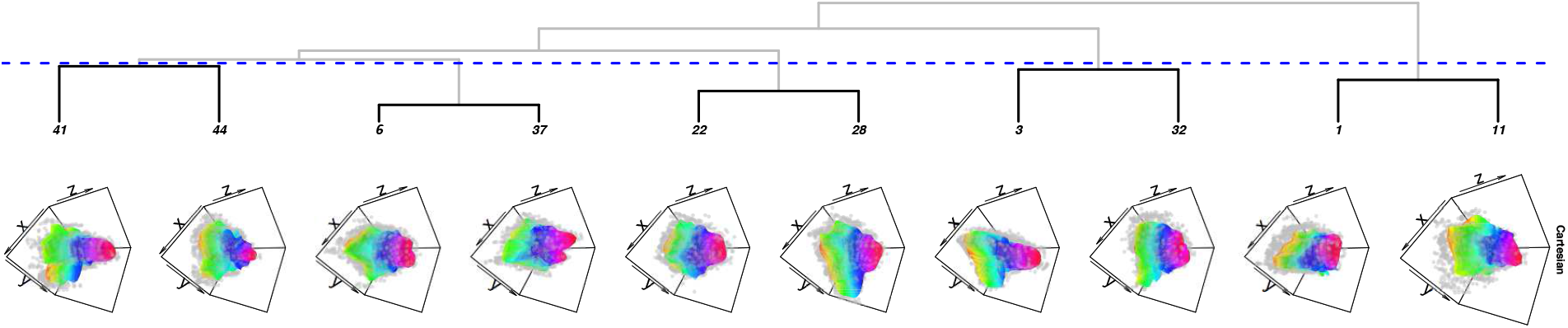
Top panel, dendrogram of the marginal likelihood associated with each cell using spherical harmonics with *L*_max_ = 12. In this example, *D* = 10, *N_i_* = 2000 for *i* = 1, 2,…, 10, *d* = (1, 3, 6, 11, 22, 28, 32, 37, 41, 44) and 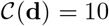. Black lines represent the improvement in the marginal likelihood and gray lines depict the deterioration in the marginal likelihood. The dashed blue line indicates the maximum a posteriori cutting point for the dendrogram, *d* = (1, 3, 6, 1, 22, 22, 3, 6, 41, 41) and 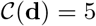. Bottom panel, the 3D data and the corresponding fit in the Cartesian coordinates.

We repeat the same procedure considering all *D* = 50 cells. The clustering result is reported in Figure 4.

**Fig 4.**
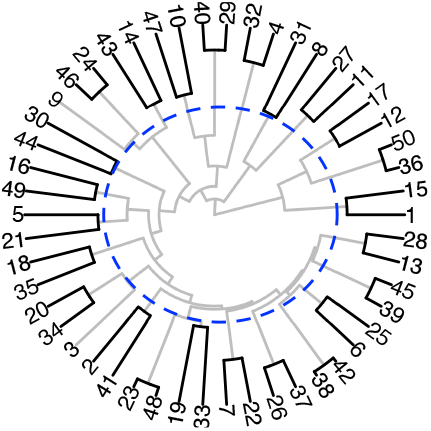
The dendrogram of the marginal likelihood associated with *D* = 50 cells using spherical harmonics with *L*_max_ = 12. Black lines represent the improvement in the marginal likelihood and gray lines depict the deterioration in the marginal likelihood. The dashed blue circle indicates the maximum a posteriori cutting point for the dendrogram.

As we discussed in Section 1.2, the number of all possible groupings is 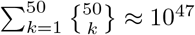. In practice, it is not feasible to explore all possible groupings when *D* is relatively large. We ran the random Gibbs sampling for 8OOO cycles as an example. The convergence behavior of the sampling throughout the 8000 cycles is reported in Figure 5.

**Fig 5.**
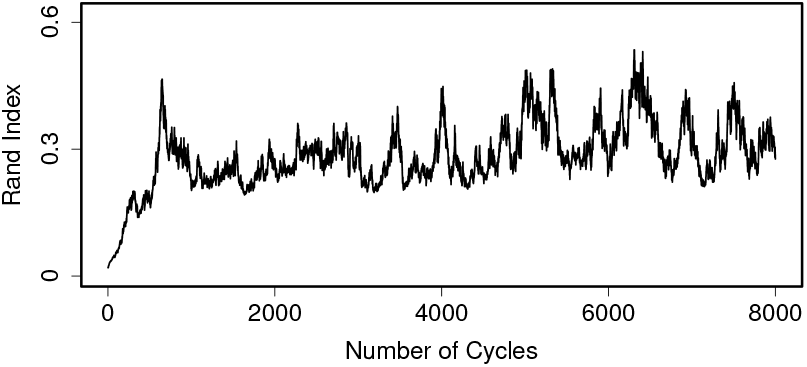
The result of random Gibbs sampling for the same cells as in Figure 4. The Rand index between the grouping suggested at each cycle of random Gibbs sampling with the grouping produced by the dendrogram in Figure 4.

The grouping generated from random Gibbs sampling after the 8000 cycles is as follows,

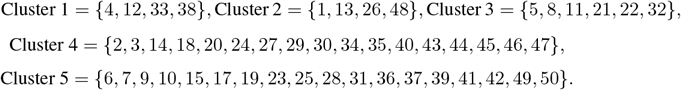

## 3 Conclusion

Having regarded the surface of a shape as a continuous function, rather than discrete landmarks, we have proposed a simple method for surface modelling of shapes such as biological cells. We have also proposed a new information criterion, called CLUSBIC, for model-based clustering and have shown that the proposed criterion is consistent.

In this work, we considered the Gaussian conjugate priors to favor computational simplicity. We proved the consistency of CLUSBIC in clustering. Note that in our settings, the increase in number of observations *N* does not necessarily imply the increase in the number of clusters 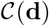 contrary to the classical clustering problem. Therefore, the consistency of CLUSBIC remains valid.

Investigation of the physical structure of cells, as simple closed shapes, can be highly useful in biology, specifically for the diagnosis of cancer. The result, in this preliminary work, shows that our proposed methodology is quite applicable and can produce promising results.

## Acknowledgments

This work was supported by the Natural Sciences and Engineering Research Council of Canada through Discovery Grants to M. Asgharian (NSERC RGPIN 217398-13).

## A

*Proof of Theorem 1*. Consider the marginal posterior for a set of *D* shapes

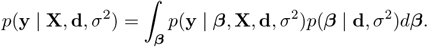

For simplicity, we use the notation *p*(**y** | *β, σ*^2^) instead of *p*(**y** | *β*, **X**, **d**, *σ*^2^). In order to obtain an approximation to this integral, we take a second order Taylor expansion of the log-likelihood at 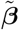, the solution to the following constrained optimization problem,

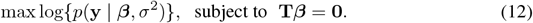

The solution 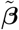 can be found using the method of Lagrange multipliers. The Lagrangian function for this problem is

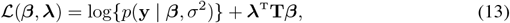

where λ is the vector of Lagrange multipliers. Expanding 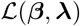 about 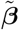 and 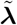.

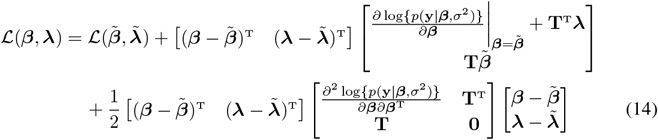

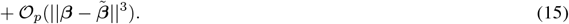

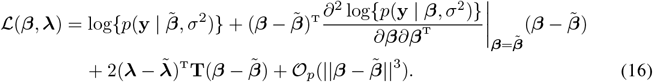

Under the assumption that **T***β* = **0**,

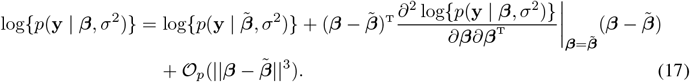

Defining the average observed Fisher information matrix as

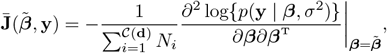

we have

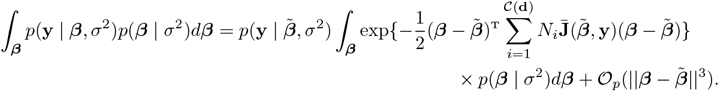

Considering 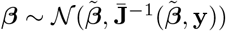 where 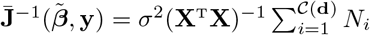,

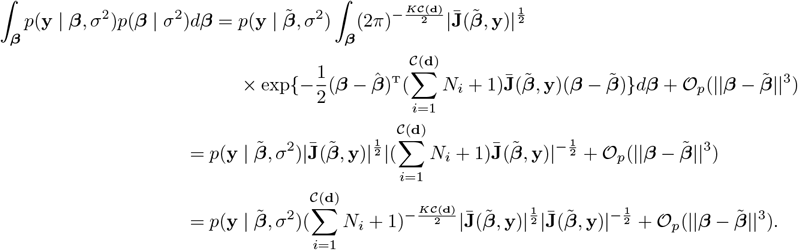

The desired result then follows upon noticing that 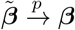 [42] and 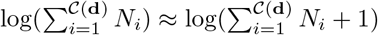 for large value of 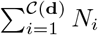.

## B

*Proof of Theorem 2*. The proof comprises two steps. First, the consistency of CLUSBIC is established for Gaussian models. We then extend the result to smooth non-Gaussian models where by “smooth “ we generally mean the likelihood is a C^3^ function of the unknown parameter. The second step essentially follows from step one upon applying a quadratic approximation to the logarithm of the likelihood.

### Step 1. Gaussian Model

Suppose the error terms, (4), are distributed according to the Gaussian distribution, one can easily show that the CLUSBIC has the following form, similar to the BIC, see [43],

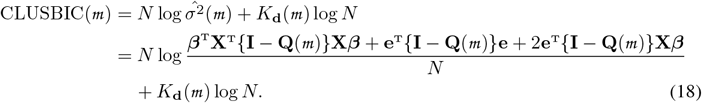

The estimate of the variance matrix for model *m* is,

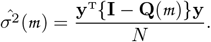

Suppose *K*_d_ and *D* are fixed, we follow the same setting as in [34] by modifying constraints of the form **T***β* = **0**. The following two assumptions are required for the proof,

1. **X**^T^**X** is positive definite.
2. 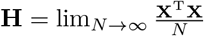 is positive definite.

The validity of these two assumptions relies on the validity of the following two assumptions for all models, i.e. ∀*i* ∈ {1, 2,…, *D*}

1. 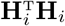 is positive definite.
2. 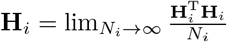 is positive definite.

#### Lemma 1.

*For* 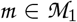, *and for any positive h*, lim_N→∞_ *N^h^p_N_*(*m*) = 0, *using* CLUSBIC.

*Proof*. The proof is given in Supplementary Material.

#### Lemma 2.

*Here, we follow a similar approach as in Lemma 1. For* 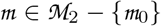, lim_*N*→∞_*p_N_*(*m*) = 0, *using* CLUSBIC.

*Proof*. The proof is given in Supplementary Material.

Having established the above lemmas, the following theorem can be established for Gaussian models.

#### Theorem 3.

*The* CLUSBIC *is a consistent clustering measure for Gaussian models*.

*Proof*. The proof is given in Supplementary Material.

### Step 2. non-Gaussian Model

First note that for a general smooth likelihood we have

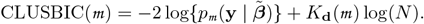

We need to show that lim_*N*→∞_ *p_N_*(*m*) = 0 for 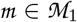 and 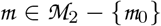 or equivalently,

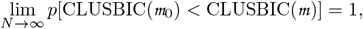

for 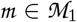 and 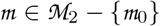. We now note that

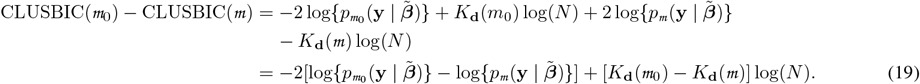

For any 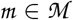 and the true value of parameter *β*_0_, one can write the following decomposition

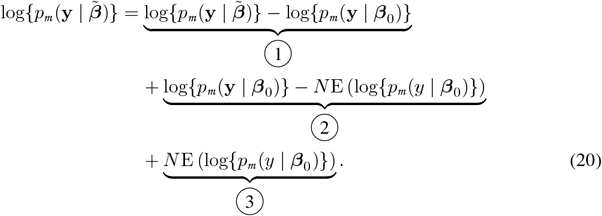

Applying the second order Taylor expansion to log{*p_m_* (**y** | *β*_0_)} at the point 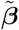, equation (17),

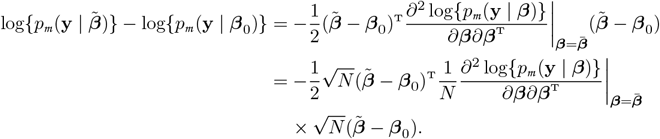

Under the usual regularity conditions, see [44, page 209]), we have

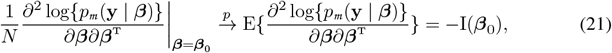

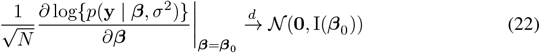

where I(*β*_0_) is the Fisher information matrix. On the other hand, by the definition of constrained optimization problem equations (12), and (13),

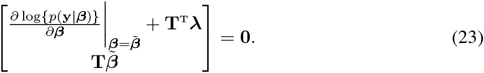

Expanding the score function around the point *β*_0_, and λ_0_ = **0**, we have

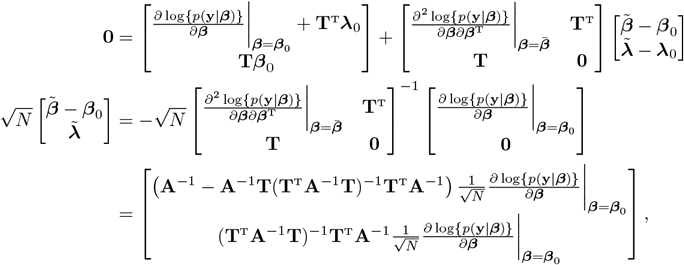

where 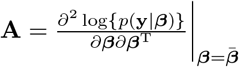. Taking into account equations (21) and (22), one can show that

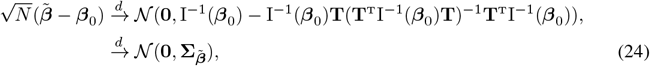

where I^-1^ (*β*_0_) is the inverse of the Fisher information matrix. By equations (24) and (21), and the fact that 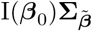 is an idempotent matrix, one can conclude that

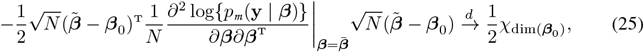

where *χ* has a chi-squared distribution with dim(**b**_0_) degrees of freedom. As the convergence in distribution implies boundedness in probability, component 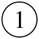 in (20) is of order 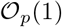.

As for component 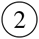 in (20), by the central limit theorem,

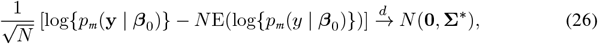

where 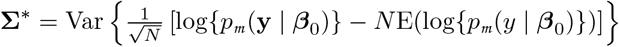. Accordingly, component 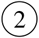 in (20) is of order 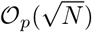.

Now coming back to the equation (19), for both 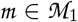 and 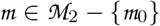,

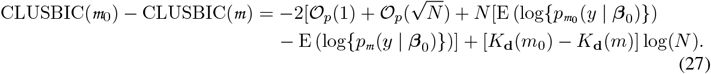

Using Jensen’s inequality,

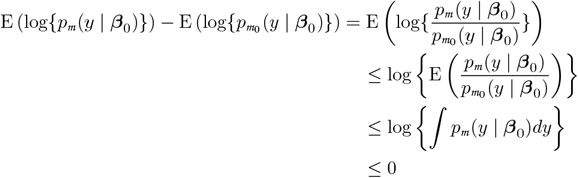

and hence

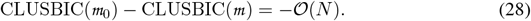

Therefore, as *N* tends to infinity

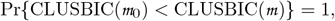

for models in both 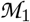 and 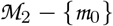. This completes the proof.

## C Supplementary Material

Here, we provide some supplementary technical materials useful in the proof of the Lemma 1, Lemma 2, and Theorem 3 in the Section B.

i. The column space of a matrix **A** is denoted by C(**A**), and defined as the space spanned by the columns of **A**.
ii. The rank of **A** is defined to be the dimension of C(**A**), dim{C(**A**)}, i,e, the number of linearly independent columns of **A**.

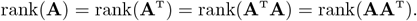
iii. Orthogonal complement of the sub-space is defined as

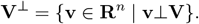
iv. If **V** ⊂ **W**, then **V**^⊥^ ⋂ **W** = {**v** ∈ **W** | **v**⊥**V**} is called the orthogonal complement of **V** with respect to **W**.

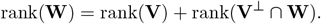
v. **Q_X_** = **X**(**X**^T^**X**)^-1^**X**^T^ is called projection matrix onto C(**X**). The matrix **Q_X_** is symmetric 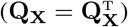 and idempotent (**Q_X_Q_X_** = **Q_X_**).
vi. If 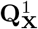 and 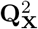 are projection matrices with 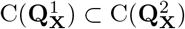, then 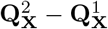 is also a projection matrix onto the orthogonal complement of 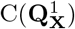 with respect to 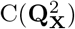.
vii. Let **A** be *k* × *k* matrix of constants and 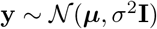. If **A** is idempotent with rank *p*, then

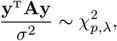

where 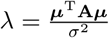.

*of Lemma 1*. In order to prove the lemma, we work with the difference between the CLUSBIC of the true model and any arbitrary model in 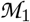. We decompose this difference into several random variables. Taking into account the properties of multivariate Gaussian distribution and the quadratic forms from the same family, we proceed with the proof. Let 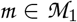,

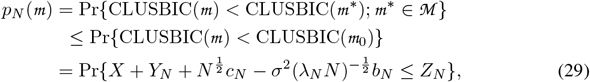

where,

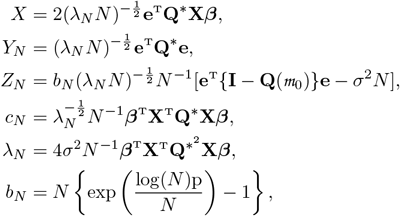

and p = *K*_**d**_(*m*_0_) – *K*_**d**_(*m*), **Q**^*^ = **Q**(*m*_0_) – **Q**(*m*). Since 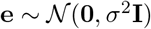, the following properties can be easily verified.

1. 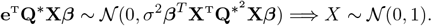
2. *Y_N_* is a quadratic form from the Gaussian distribution.
3. Since {**I** – **Q**(*m*_0_)} is a symmetric, idempotent matrix,

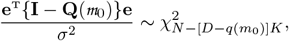

where 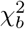 is the chi-squared distribution with *b* degrees of freedom.

The validity of equation (29) can be easily verified through the definition of CLUSBIC in equation (18). By the Fréchet inequality,

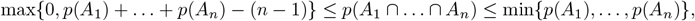

where *A_i_*’s are some events, the equation (29) is bounded by,

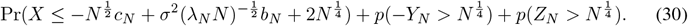

Using the assumptions 1 and 2,

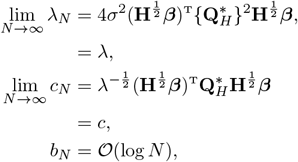

where 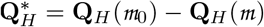, and

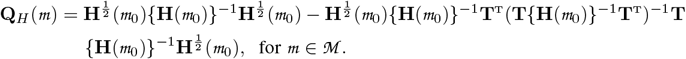

Let 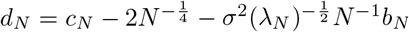. One can easily show that 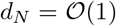 as lim_*N*→∞_ *c_N_* = *c* and 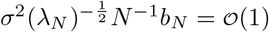. Using the characteristics of the standard Gaussian distribution function,

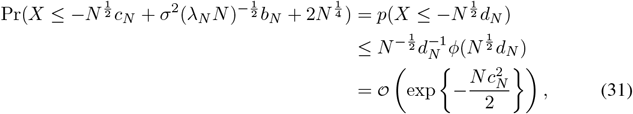

where *ϕ*(.) is the density function of the standard Gaussian distribution.

Given that *Y_N_* is a quadratic form, using the definition of moment generating functions for quadratic forms [45], we have

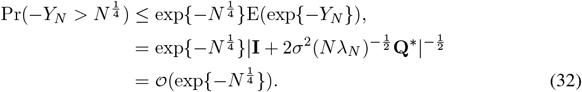

By the property 3,

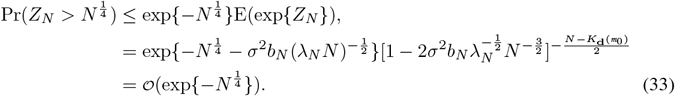

The equations (31), (32) and (33) complete the proof.

*of Lemma 2*. For 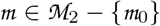,

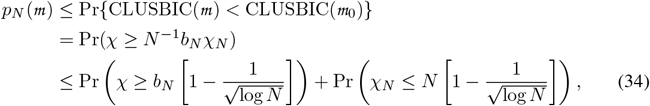

where

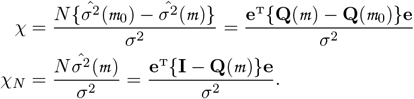

By definition, we have

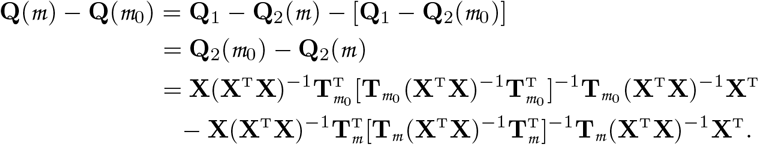

Under *H*_0_, for an arbitrary model we have, **T***β* = **0**,

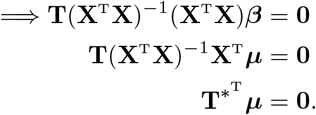

Under *H*_0_, *μ* ∈ C(**X**) = **V** and *μ*⊥C(**T**\*), or *μ* ∈ C(**T**\*)^⊥^ ⋂ C(**X**) = **V**_0_ which is the orthogonal complement of C(**T**\*) with respect to C(**X**).

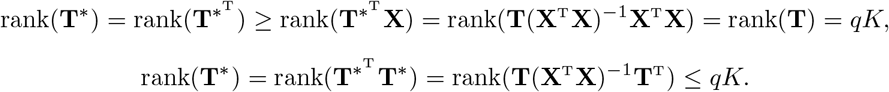

Therefore, rank(**T**\*) = *qK*. By definition of projection matrix (v),

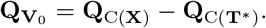

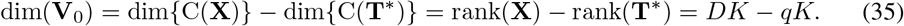

By the property (vi), **Q**(*m*) – **Q**(*m*_0_) is a projection matrix. Thus, using equation (35), *χ* has chi-squared distribution with p degrees of freedom,

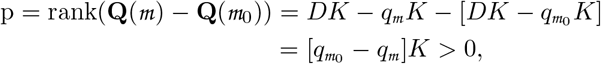

since *q*_*m*_0__ > *q_m_*. Similarly, *χ_N_* has chi-squared distribution with *N* – [*DK* – *q_m_K*] degrees of freedom.

Back to equation (34), since 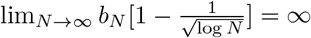,

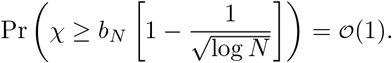

For the second term in equation (34), one can show that

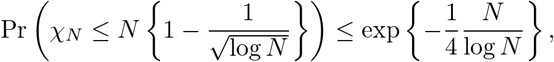

using an inequality on chi-squared distribution, see [46]. This completes the proof.

*of Theorem B 3*. The equation (11) follows from Lemma 1 and Lemma 2.

The risk, or expected loss, for the model is

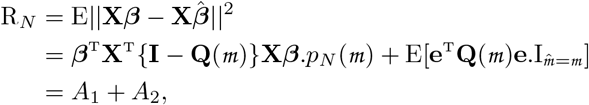

where 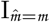 is the indicator function. By the Cauchy-Schwartz’s inequality,

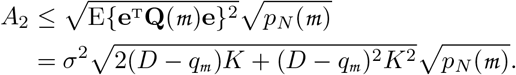

For 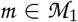

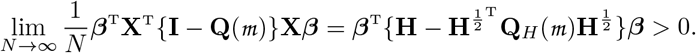

Equivalently,

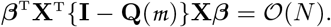

Therefore, *A*_1_ and *A*_2_ tend to 0 as *N* → ∞ by condition (a).

For 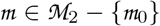, *A*_2_ → 0 as *N* → ∞ by condition (b). In addition,

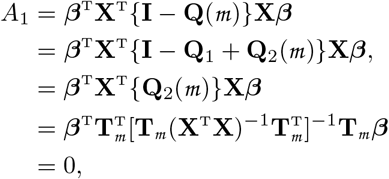

since **T**_*m*_ = **0** for models in 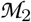.

Consequently, lim_*N*→∞_ R_*N*_ = 0 for both 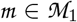 and 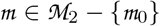. Therefore, if conditions (a) and (b) are satisfied,

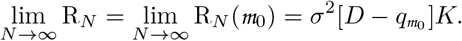

This completes the proof of Theorem.

## Footnotes

1 http://murphylab.web.cmu.edu/data/#3DHeLa

